# Caudal *Fgfr1* disruption produces localised spinal mis-patterning and a terminal myelocystocele-like phenotype in mice

**DOI:** 10.1101/2023.03.07.531511

**Authors:** Eirini Maniou, Faduma Farah, Zoe Crane-Smith, Andrea Krstevski, Athanasia Stathopoulou, Nicholas D.E. Greene, Andrew J. Copp, Gabriel L. Galea

## Abstract

Closed spinal dysraphisms are poorly understood neurodevelopmental malformations commonly classed as neural tube defects. Several, including terminal myelocystocele, selectively affect the distal lumbosacral spine. We previously identified a neural tube closure-initiating point, Closure 5, involved in forming the distal spine of mice. Here we document equivalent morphology of the caudal-most end of the closing posterior neuropore (PNP) in mice and humans, suggesting Closure 5 is conserved in humans. It forms in a region of active fibroblast growth factor (FGF) signalling and pharmacological blockade of FGF receptors (Fgfr) impairs Closure 5 formation in cultured mouse embryos. Conditional genetic deletion of *Fgfr1* in caudal embryonic tissues with *Cdx2*^*Cre*^ similarly impairs Closure 5 formation and leads to morphologically abnormal PNPs, which nonetheless achieve delayed closure although delayed. After PNP closure, a localised region of the distal neural tube of *Fgfr1*-disrupted embryos re-opens into a trumpet-like flared central canal between the presumptive hindlimbs, progressing to form a distal fluid-filled sac overlying ventrally flattened spinal cord. This phenotype resembles terminal myelocystocele. Histological analysis of spinal progenitor domains reveals regional and progressive loss of ventral spinal cord progenitor domains preceding cystic dilation of the central canal. Initially, the Shh and FoxA2-positive ventral domains are lost, resulting in Olig2-labelling of the ventral-most neural tube. The Olig2-domain is also subsequently lost, eventually producing a neural tube entirely positive for the dorsal marker Pax3. Thus, a terminal myelocystocele-like phenotype can arise after completion of neural tube closure due to localised spinal mis-patterning caused by disruption of Fgfr1 signalling.

## Introduction

Neural tube defects (NTDs) are a heterogenous group of congenital malformations resulting from dysmorphogenesis of the neural tube (NT), the embryonic precursor of the central nervous system. The most common and severe NTDs are those caused by failure to close the NT, including open spina bifida (myelomeningocele). Successful closure of the NT during primary neurulation requires coordinated deformation of the neural plate into a continuous tube of neuroepithelial cells covered by surface ectoderm along the back of the embryo (1). Closure begins at distinct closure-initiating points where the opposite halves of the embryo are first brought into apposition at the dorsal midline. Five closure-initiating points had originally been proposed to initiate NT closure, based on anatomical clustering patterns of NTD lesions in human embryos (2, 3). Three closure-initiating points were subsequently experimentally identified in mouse embryos: Closure 1 at the hindbrain/cervical boundary, Closure 2 at the forebrain/midbrain boundary (absent in humans) and Closure 3 at the ventral forebrain (4). The regions of open NT between these Closure points, called neuropores, close through progressive zippering (1). No closure-initiating point is present where Closure 4 was initially inferred, but we have more recently confirmed the presence of Closure 5 at end of the presumptive spinal NT in mice (5, 6). Closure 5 is a load-bearing tissue structure which physically holds the neural folds together at the dorsal midline during an approximately 10-hour period from when it is first morphologically identifiable (∼25 somite stage) until the spinal NT is fully closed (∼30 somite stage). Closure 5 was initially predicted in humans due to clusters of very distal spina bifida (2), but this ephemeral structure has not yet been visualised in human embryos.

In mice, surface ectoderm cells at Closure 5 extend cellular ruffles, consistent with zippering in a caudal-to-cranial direction (5). Thus, the presumptive spinal neuropore, called the posterior neuropore (PNP), closes primary due to caudally-directed zippering from Closure 1, biomechanically aided at late stages by Closure 5. The NT lumen extends caudally beyond Closure 5 through a process of mesenchyme-to-epithelium transition with lumen formation referred to as secondary neurulation, forming the sacral and coccygeal spine. Several important but poorly understood NTDs characteristically localise to the junctional region between primary and secondary neurulation, where Closure 5 forms (7, 8). These include terminal myelocystocele, often described as a “closed” form of spina bifida because the dysmorphic spinal cord is covered with skin (9). Terminal myelocystocele characteristically presents as a trumpet-like flaring of the spinal cord’s distal central canal, forming a localised cystic dilation at the terminal end of the body which continues to expand postnatally (9, 10). It is commonly associated with other malformations, including of musculoskeletal and urogenital structures, and variable neurological dysfunction (10).

There are, to our knowledge, no established animal models of terminal myelocystocele which mechanistically explain its pathogenesis. The most widely-cited hypothesis for its formation relates to a balloon-like dilation observed at the junction between the primary and secondary NT of chick embryos (9), although this is an aspect of normal neurulation and is not a pathological finding. The joined primary and secondary NT form a continuous lumen bordered by neuroepithelial cells which differentiate into dorsoventrally patterned neural lineages. Many progenitor domains are similar between the primary and secondary NT, including a dorsal Pax3-expressing domain, intermediate Pax6 domain and ventral FoxA2 domain (11). However, there are regional differences in neural patterning including the absence of motor neuron differentiation in the caudal NT region formed through secondary neurulation (11). Motor neuron progenitors (pMN) differentiate in the ventral primary neural tube and are characteristically identified by expression of the transcription factor Olig2, promoted by high levels of Sonic hedgehog (Shh) morphogen from the NT floorplate and underlying notochord (12, 13). Shh is the best-established inducer of ventral NT fates (14, 15), but does not act in isolation. For example, fibroblast growth factor (FGF) ligands promote maintenance of spinal neural progenitor identities while restricting differentiation into post-mitotic neurons (16-18). FGF signalling opposes somitic mesoderm-derived retinoic acid, which promotes neuron terminal differentiation (19, 20). Loss of the retinoic acid synthesising enzyme *Raldh2* does not prevent neuroepithelial commitment, as shown by Sox2 expression, but impairs differentiation of dorsoventral progenitor domains including ventral Olig2 and intermediate Pax6 in mice (21).

Insults which change the timing of neural progenitor differentiation have also been associated with open spina bifida in mice, such as loss of the dorsal neural crest inducer Pax3 (22, 23) or unrepressed Shh signalling causing expansion of ventral neural tube fates (24, 25). Studies into the interplay between neurulation and subsequent neurogenesis have been limited by the severely abnormal morphology of embryos lacking key mediators. For example, embryos globally lacking the FGF receptor Fgfr1 die soon after gastrulation without closing any portion of their NT (26). FGF pathway components including ligands such as neuroepithelial Fgf8 are particularly enriched in the caudal-most part of embryo, where Closure 5 forms (27).

Here we initially used a combination of pharmacological antagonism and regional genetic deletion to test the contributions of FGF signalling, specifically through Fgfr1, in PNP closure and Closure 5 formation. Conditional genetic deletion of *Fgfr1* produces mouse fetuses which are viable until birth but have a cystic dilation of the distal spinal NT forming a closed spinal dysraphism. We find that the embryological origins of this malformation involve regional and progressive loss of NT ventral progenitor domains prior to the closed and dorsalised NT flattening ventrally while retaining its dorsal skin covering, closely resembling terminal myelocystocele in humans.

## Results

### Gradual neural fold elevation forms Closure 5 at late stages of spinal closure

Human primary neurulation is completed by Carnegie Stage (CS)13, around 30-35 days of gestation (28). At CS11, the PNP is open and has a spade-like morphology (Figure 1A) equivalent to the mouse PNP on embryonic day E9.5 (Figure 1B). Progression of mouse PNP closure involves caudal narrowing to form an elliptical opening surrounded by purse string-like F-actin cables as Closure 5 forms at the PNP’s caudal extreme (Figure 1B). Although late-stage human PNPs are not available to us for 3D imaging, brightfield images archived by the Human Developmental Biology Resource suggest that equivalent changes in shape to form an elliptical PNP with caudal Closure 5 in humans as in mice (Supplementary Figure 1).

**Figure 1:**
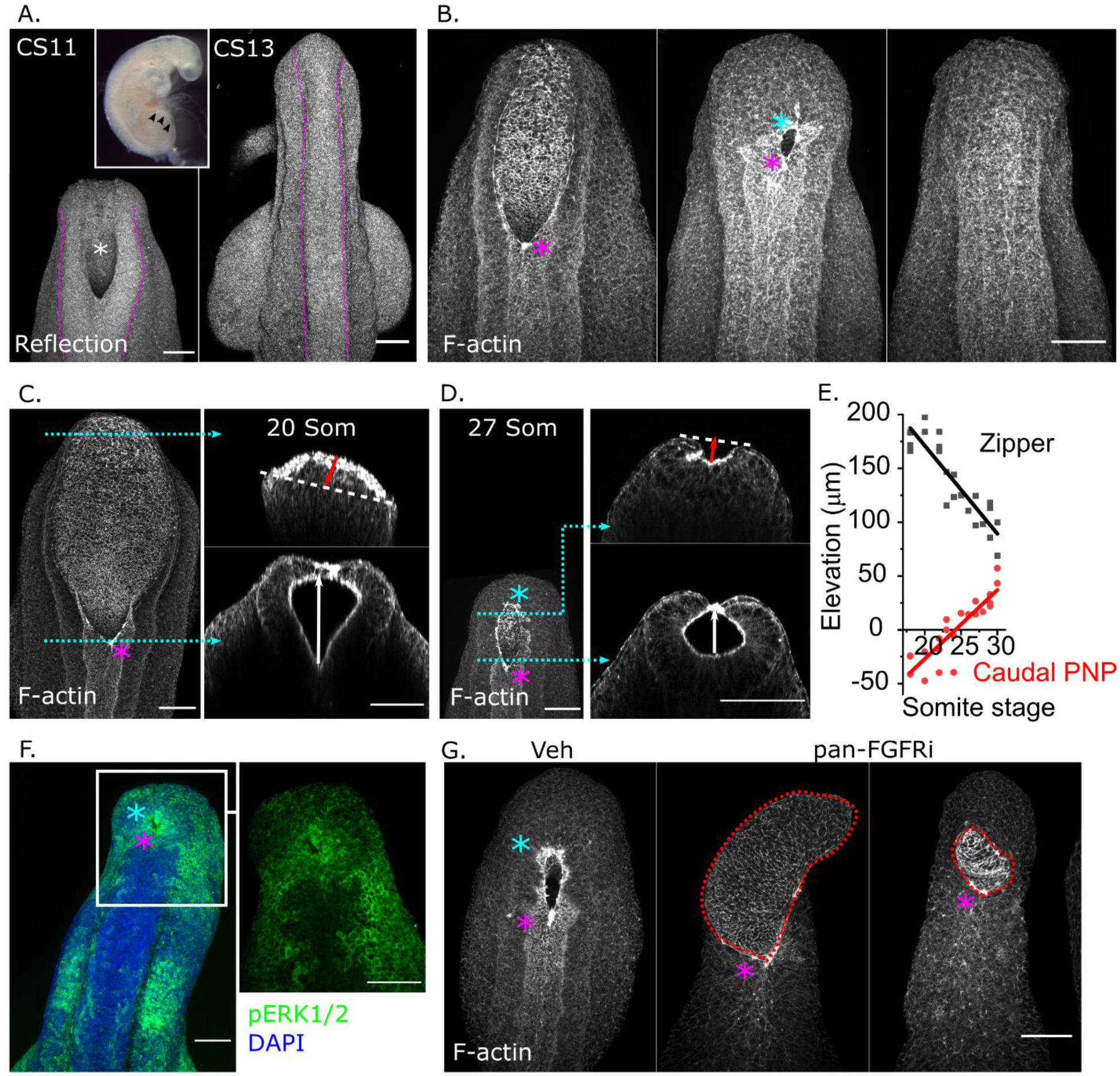
FGF-dependent elevation of the distal neural folds forms Closure 5. A. Reflection confocal images of the PNP of a CS11 (* and black arrowheads in insert) and closed neural tube of a CS13 human embryo. Dashed magenta lines indicate the borders of the neuroepithelium. Scale bars = 50 μm. B. Representative confocal images of phalloidin-stained mouse PNPs at sequential stages of closure (E9.5-E10.5) illustrating the formation of Closure 5 and completion of closure. Scale bar = 100 μm. C. Confocal image of a phalloidin-stained PNP from a 20-somite stage mouse embryo showing optical cross-sections at the level of the zippering point and at 90% of the PNP’s length from the proximal end. The white arrow indicates zippering point elevation, red arrow indicates eversion of the caudal neural folds relative to the apical surface of the neuroepithelium. D. Equivalent to C showing a 27-somite stage mouse embryo PNP indicating elevation of the caudal neural folds. E. Quantification of elevation at the zippering point and caudal neural folds (90% of the PNP’s length) in mouse embryos at the indicated somite stages. F. Wholemount-stained confocal image of a 29 somite-stage mouse PNP showing pERK1/2 immunolocalization. Scale bar = 100 μm. G. Confocal images of a vehicle-treated and two pan-FGFR inhibitor-treated mouse embryos after 24 hours of whole embryo culture. Dashed red lines indicate the abnormal neuroepithelium which protrudes out of the PNP in inhibitor-treated embryos. Scale bar = 100 μm. Magenta asterisks indicates the rostral zippering point, cyan asterisks indicate Closure 5.

We previously demonstrated that Closure 5 is a load-bearing structure (5, 6). Here we observe that it forms through gradual elevation of the caudal neural fold tips. The caudal PNP neuroepithelium is everted at earlier developmental stages, but progressively becomes apically concave as the neural fold tips elevate dorsally in embryos with >25 somites (Figure 1C-E), when Closure 5 typically forms. Dorsal elevation of the rostral zippering point follows an inverse pattern: it is initially elevated ∼200 μm above the ventral-most apical neuroepithelium, but its elevation halves to ∼100 μm in later-stage embryos with established Closure 5 (Figure 1E). The continuum of neural fold morphologies, from early eversion to late elevation, suggests coordinated morphogenesis of the caudal PNP to complete primary neurulation.

### Pharmacological blockade of FGF signalling diminishes Closure 5 formation

The caudal PNP is known to be enriched in components of cascades including canonical Wnt and Fgf signalling (Supplementary Figure 2). Fgf components are particularly prominent (Supplementary Figure 2) and we observe marked phosphorylation of the pathway’s putative down-steam effector ERK1/2 selectively around Closure 5 (Figure 1F). Pharmacological inhibition of Fgfr signalling in mouse whole embryo culture with BGJ 398 does not change somite gain (vehicle 27±3 somites, pan-Fgfr inhibitor 26±3 somites at the end of culture, mean±SD), but produces highly irregular PNP morphologies, particularly of the caudal-most PNP where abnormal tissue outgrowth forms (Figure 1G). The range of morphologies observed in Fgfr-inhibited embryos suggests abnormal progression of closure (Supplementary Figure 3A). Cells within this outgrowth are derived from neuromesodermal progenitors lineage-traced with *T*^*CreERT2*^, which continues to lineage-trace both neural tube and mesoderm cells after 24 hours of Fgfr inhibition (Supplementary Figure 3B). Cells in the PNP outgrowth also continue to express the neuroepithelial marker N-cadherin (Supplementary Figure 3C).

Phosphorylation of ERK1/2 around the PNP rim persists despite Fgfr inhibition (Figure 2A). Expression of the pathway ligand *Fgf8* is retained or increased in the PNP neuroepithelium, whereas it is abolished in the forelimb bud of Fgfr-inhibited embryos (Figure 2B), indicating tissue-specific regulation. Retinoic acid signalling is known to be antagonised by caudally-active Fgf signalling (29). Pharmacological Fgfr blockage causes caudal expansion of the retinoic acid signalling domain genetically labelled by a retinoic acid response element (RARE) driving LacZ expression (Figure 2C-D). However, RARE activity does not detectably extend to the caudal-most PNP region where Closure 5 should form (Figure 2C). This caudal PNP region fails to elevate, remaining everted, in contrast to vehicle-treated controls which elevate their caudal PNP beyond the 25 somite stage (Supplementary Figure 3E). Fgfr inhibitor-treated embryos largely fail to form Closure 5 and do not complete PNP closure by E10.5 (Supplementary Figure 3F).

**Figure 2:**
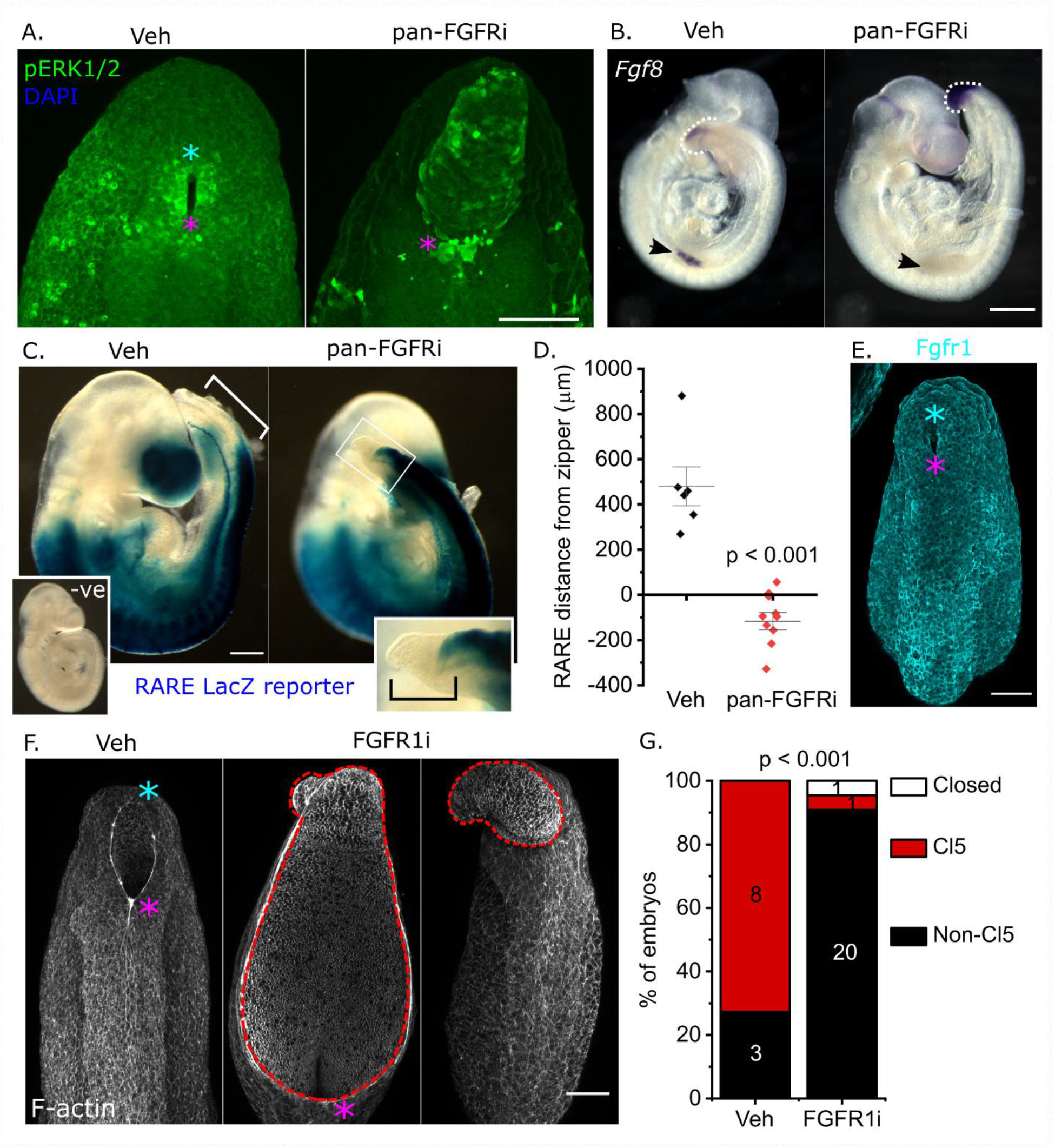
Tissue-specific functions for FGF signaling, likely through Fgfr1, promote timely completion of PNP closure. A. Wholemount-stained confocal images showing pERK1/2 immunolocalization in a vehicle and pan-FGFRi-treated embryos after 24 hours of whole embryo culture. Note persistence of pERK-bright cells around the PNP rim in the pan-FGFRi-treated embryo. Scale bar = 100 μm. B. Wholemount *Fgf8* in situ hybridisation in vehicle- and pan-FGFRi-treated embryos after 8 hours of whole embryo culture. Dashed white lines indicate the end of the tailbud, arrows indicate the forelimb buds. Scale bar = 500 μm. C. Brightfield images showing RARE-mediated LacZ expression domain in vehicle and pan-FGFRi-treated embryos after 24 hours of whole embryos culture. An equivalently processed negative (-ve) control embryo lacking LacZ is also shown. The double-headed white arrow indicates the distance between the RARE domain and the zippering point. The black bracket in the inset indicates that the RARE domain does not extend to the end of the PNP in pan-FGFRi-treated embryos. Scale bar = 350 μm. D. Quantification of RARE domain distance to the end of zippering point in vehicle and pan-FGFRi-treated embryos after 24 hours of whole embryo culture. E. Wholemount immunolocalization of Fgfr1 in a 28-somite stage embryo. Scale bar = 100 μm. F. Confocal images of embryos treated with vehicle or a second, Fgfr1-targeting antagonist after 24 hours of whole embryo culture. Dashed red lines outline the abnormal neuroepithelium in inhibitor-treated embryos. Scale bar = 100 μm. G. Quantification of the proportion of embryos with closed PNPs, open PNPs with Closure 5 morphology (caudally narrowed PNP with elevated caudal neural folds which meet dorsally) or open PNPs without Closure 5 in vehicle and FGFR1-targeting inhibitor-treated embryos after 24 hours of whole embryo culture. Numbers indicate the number of embryos observed in each category. Magenta asterisks indicates the rostral zippering point, cyan asterisks indicate Closure 5.

Various Fgf receptors are known to be expressed in and around the PNP and may have both redundant and obligate roles. *Fgfr3* knockout causes post-natal phenotypes without neural tube defects (30). *Fgfr2* embryos die ∼E11.5, after successful closure of the neural tube (31). Fgfr1 is expressed diffusely around the PNP rim (Figure 2E) (32) and its chimeric deletion causes distal spina bifida (33). We therefore focused on *Fgfr1* as the more likely non-redundant mediator of FGF signalling promoting PNP closure. Treating cultured embryos with a second, Fgfr1-targeting pharmacological antagonist PD 173074 (34, 35) also did not change somite gain (vehicle 28±3 somites, pan-Fgfr inhibitor 29±2 somites at the end of culture, mean±SD) and similarly caused neuroepithelial outgrowth, failure of caudal elevation and failure of Closure 5 formation (Figure 2F-G).

### *Fgfr1* conditional deletion prevents Closure 5 formation, but does not stop spinal closure

To avoid early lethality seen in global knockouts, we used a conditional approach to eliminate *Fgfr1* expression after initiation of neural tube closure with paternally-inherited CDX2P-NLS Cre (36) (henceforth *Cdx2*^*Cre*^) driving deletion of a previously-reported (37) *Fgfr1* conditional allele (*Fgfr1*^*Fl/Fl*^). *Cdx2*^*Cre*^ recombines in all embryonic tissues caudal to the cervical spine with some notable exceptions: at E10.5 it does not recombine in the notochord or lateral plate mesoderm (Figure 3A-B). Heterozygous *Cdx2*^*Cre/+*^*Fgfr1*^*Fl/+*^ (henceforth Cre;Fl/+) mice are viable and fertile. Embryos with this genotype are morphologically indistinguishable from their Cre-negative littermates and are therefore included as ‘controls’. Homozygous *Cdx2*^*Cre/+*^*Fgfr1*^*Fl/Fl*^ (henceforth Cre;Fl/Fl) embryos appear caudally truncated, with an open PNP at E10.5 (Figure 3A). Loss of *Fgfr1* does not impair embryo growth between the forelimb and hindlimb buds, but selectively truncates caudal elongation from the hind limbs to the end of the body (Figure 3B). At this stage their PNP neuroepithelium is irregularly shaped with corrugated neuroepithelium (Figure 3C-D), reminiscent of Fgfr antagonist-treated embryos in culture.

**Figure 3:**
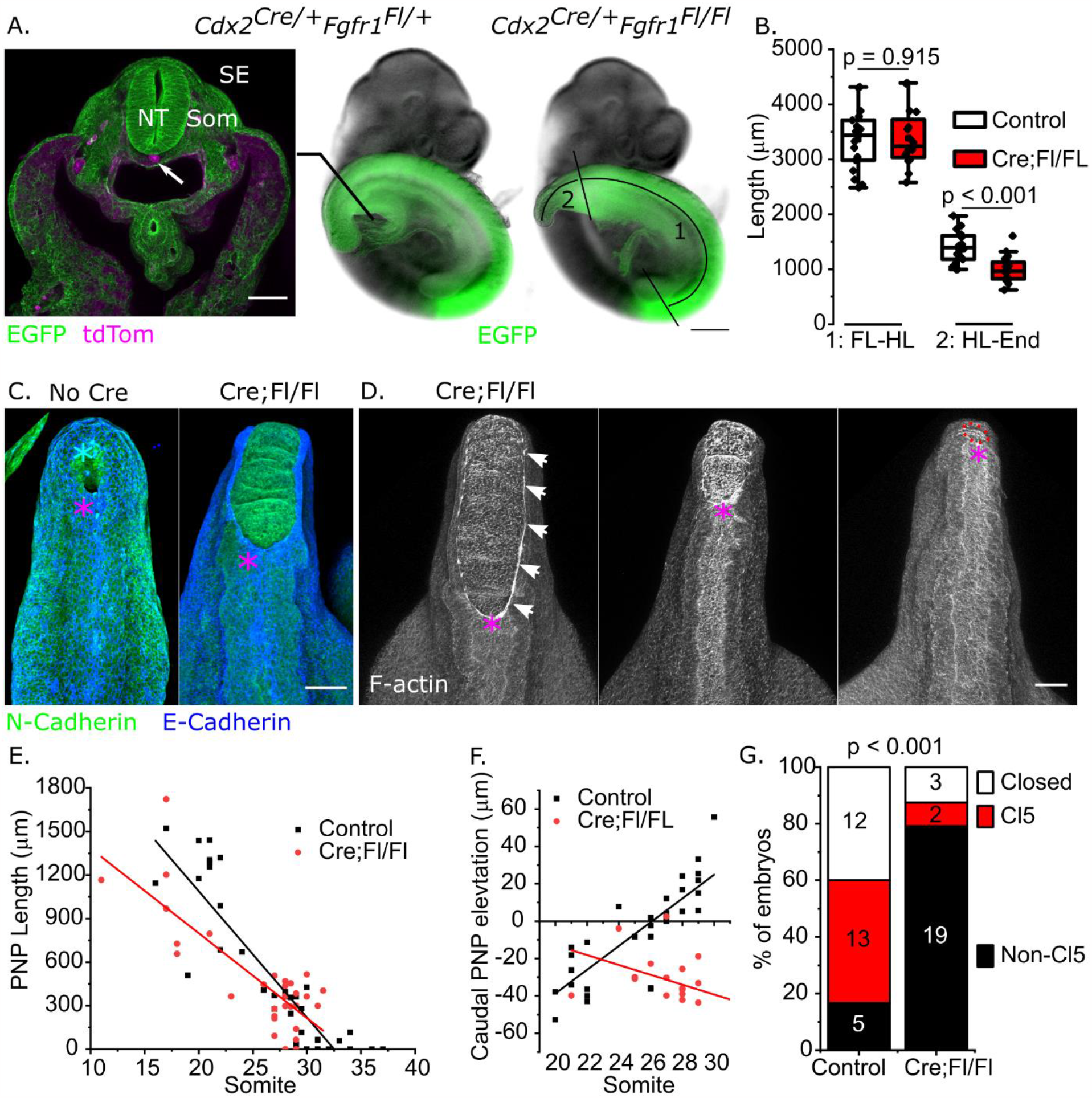
Caudal genetic deletion of Fgfr1 impairs Closure 5 formation and progression of PNP closure without diminishing embryo viability. A. E10.5 embryos showing the *Cdx2*^*Cre*^-recombination domain lineage traced using mTmG (EGFP) as a confocal-imaged cross-section and brightfield images. NT neural tube, Som somite, SE surface ectoderm. 1: forelimb (FL) to hindlimb (HL) distance, 2: hindlimb to tail end distances quantified in B. Scale bars = 100 μm (confocal) and 400 μm (brightfield). B. Quantifications of FL-HL and HL-tail end lengths in 28-31 somite stage embryos. C. Confocal images of wholemount immunoflourecently stained embryos showing E- and N-cadherin localization in E10.5 embryos with no Cre and Cre;Fl/Fl littermate. D. Wholemount images of three phalloidin-stained Cre;Fl/Fl embryos collected at E10.5 suggesting continued PNP closure. Scale bars = 100 μm. E. Quantification of PNP length versus somite stage in embryos collected at E9.5-E10.5. F. Quantification of caudal neural fold elevation (90% of PNP length) in in embryos collected at E9.5-E10.5. G. Quantification of the proportion of embryos collected at E10.5 with closed PNPs, open PNPs with Closure 5 morphology or open PNPs without Closure 5. Numbers indicate the number of embryos observed in each category. Magenta asterisks indicates the rostral zippering point, cyan asterisks indicate Closure 5.

Cre;Fl/Fl embryos continue to assemble long F-actin cables bordering the PNP as seen in wild-type embryos, and achieve progressive shortening of the PNP (Figure 3D,E). The caudal PNP elevates progressively in control embryos, whereas in Cre;Fl/Fl it tends to become increasingly everted (Figure 3F). Consequently, in embryos which had >25 somites, only 8.3% of Cre;Fl/Fl embryos 43% of control embryos have a PNP morphology indicative of Closure 5, compared with 43% of control embryos only 8.3% of Cre;Fl/Fl embryos (Figure 3G). Thus, caudal deletion of Fgfr1 causes morphologically abnormal PNPs which typically lack Closure 5 formation.

### *Fgfr1* disruption causes a terminal myelocystocele-like phenotype

A significantly smaller proportion of Cre:Fl/Fl than control embryos achieve PNP closure by E10.5 and a sub-set remain open at E11.5 (Figure 4A), presenting open lesions caudal to the hind limb buds (Supplementary Figure 4), whereas all control embryos close their PNP by this timepoint (Figure 4B). All fetuses collected from E12.5 onwards had closed neural tubes (Figure 4A), defined as surface ectoderm/skin fully covering the developing spinal cord. Control embryos at E11.5 have a closed quasi-cylindrical neural tube extending past paired somites into the tail bud (Figure 4B). Somites flanking the closed neural tube within the embryonic tail are markedly hypoplastic in Cre;Fl/Fl embryos (Figure 4B).

**Figure 4:**
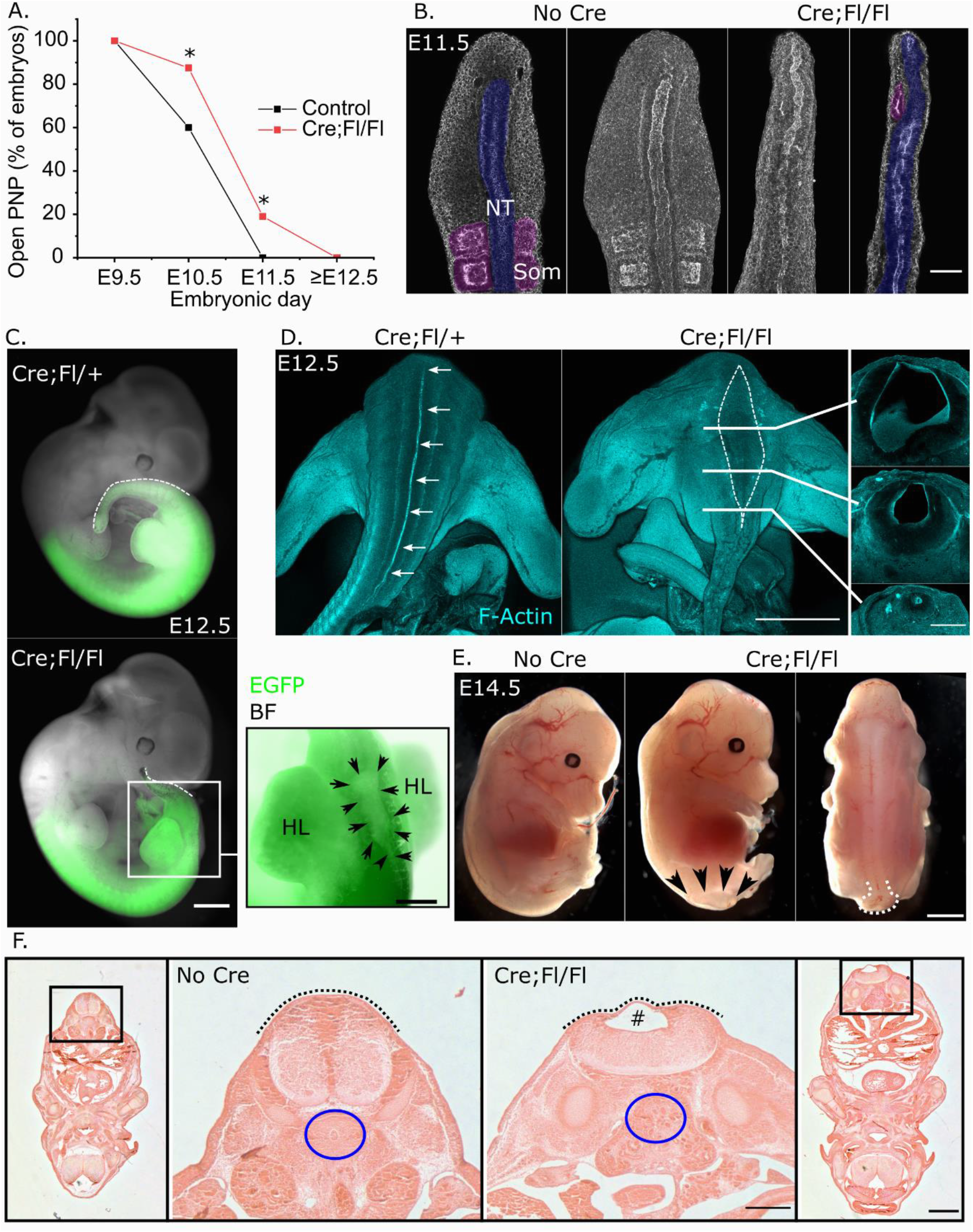
Regional deletion of Fgfr1 causes delayed PNP closure and localized terminal dilation of the neural tube central canal. A. Proportion of control and Cre;Fl/Fl embryos collected with open spinal neural tubes at the days indicated. * p < 0.05 by Fisher’s exact test. B. Wholemount maximum projections flanked by optical cross-sections through the caudal end of a control and Cre;Fl/Fl littermate showing the neural tube (NT) and adjacent somites (Som). Scale bar = 100 μm. C. Brightfield images showing the *Cdx2*^*Cre*^ recombination domain (green, EGFP) at E12.5 in a Cre;Fl/+ control and Cre;Fl/Fl littermate. The dashed white line indicates the dorsal tail border. Black arrowheads in the inset demarcate the cystic neural tube lumen. Scale bars = 300 μm (whole embryo) and 150 μm (inset). BF brightfield. D. Wholemount phalloidin stained embryos showing the continuous, narrow neural tube lumen in the control (arrows) in the control and dilated lumen (dashed white line) in the Cre;Fl/Fl embryo. Solid lines indicate the positions of sections taken through the same Cre;Fl/Fl embryo. Scale bars = 1 mm (wholemount) and 250 μm (section). E. Brightfield images of a control and littermate Cre;Fl/Fl fetuses collected at E14.5. Black arrows indicate the extent of the spinal lesion in the Cre;Fl/Fl embryo and the white dashed line indicates its sac-like dilation. Scale bar = 750 μm. F. Hematoxylin and eosin-stained sections through control and Cre;Fl/Fl littermates collected at E14.5. Dashed black lines indicate continuous skin covering, # indicates cystic dilation of the spinal cord lumen, blue circles indicate the ventral vertebral body in the control which is absent in the Cre;F/Fl. Scale bars = 300 μm (low-mag) and 100 μm (high-mag).

At E12.5, Cre:Fl/Fl embryos develop as cystic dilation of the neural tube lumen selectively between the hindlimbs (Figure 4C-D). Serial cross-sections reveal a trumpet-like flaring of the central canal with a thin dorsal neuroepithelium overlying the dilated portion (Figure 4D). By E14.5 the dorsal thinning of the roofplate and canal dilation progresses to form a ventrally-flattened spinal cord which has an ‘open’ morphology, yet is covered by skin overlying a fluid-filled sac (Figure 4E-F). This sac-like lesion at the end of the spinal cord morphologically resembles terminal myelocystocele lesions. Additional malformations are evident in these fetuses, including absence of the ventral vertebral body (Figure 4F), limb malformations and apparent delay in perineal fusion (Supplementary Figure 5).

We immunolabelled neural progenitors to confirm the identity of cells in the misshapen spinal cord of Cre;Fl/Fl embryos. Sox2 labels a continuous neuroepithelial layer encircling the central canal, but this layer is thinned dorsally in Cre;Fl/Fl embryos at E14.5 (Figure 5A). This dorsal neuroepithelium lies ventral to the overlying skin layer which stains positive for E-cadherin (Figure 5A). This dysmorphology arises in a region of closed neural tube between the hindlimb buds which is morphologically normal at E11.5 (Figure 5B). At this stage, Tuj1 positive neurons are specified in both control and Cre;Fl/Fl embryos (Figure 5B). Dorsal views of the neural tube show distinct parallel populations of Tuj1 positive neurons emerging between the hind limb buds in both, but more prominently in Cre;Fl/Fl than control embryos (Figure 5C). Consequently, the Tuj1-stained area, relative to the area of visible neural tube, is greater in Cre;Fl/Fl embryos (Figure 5D).

**Figure 5:**
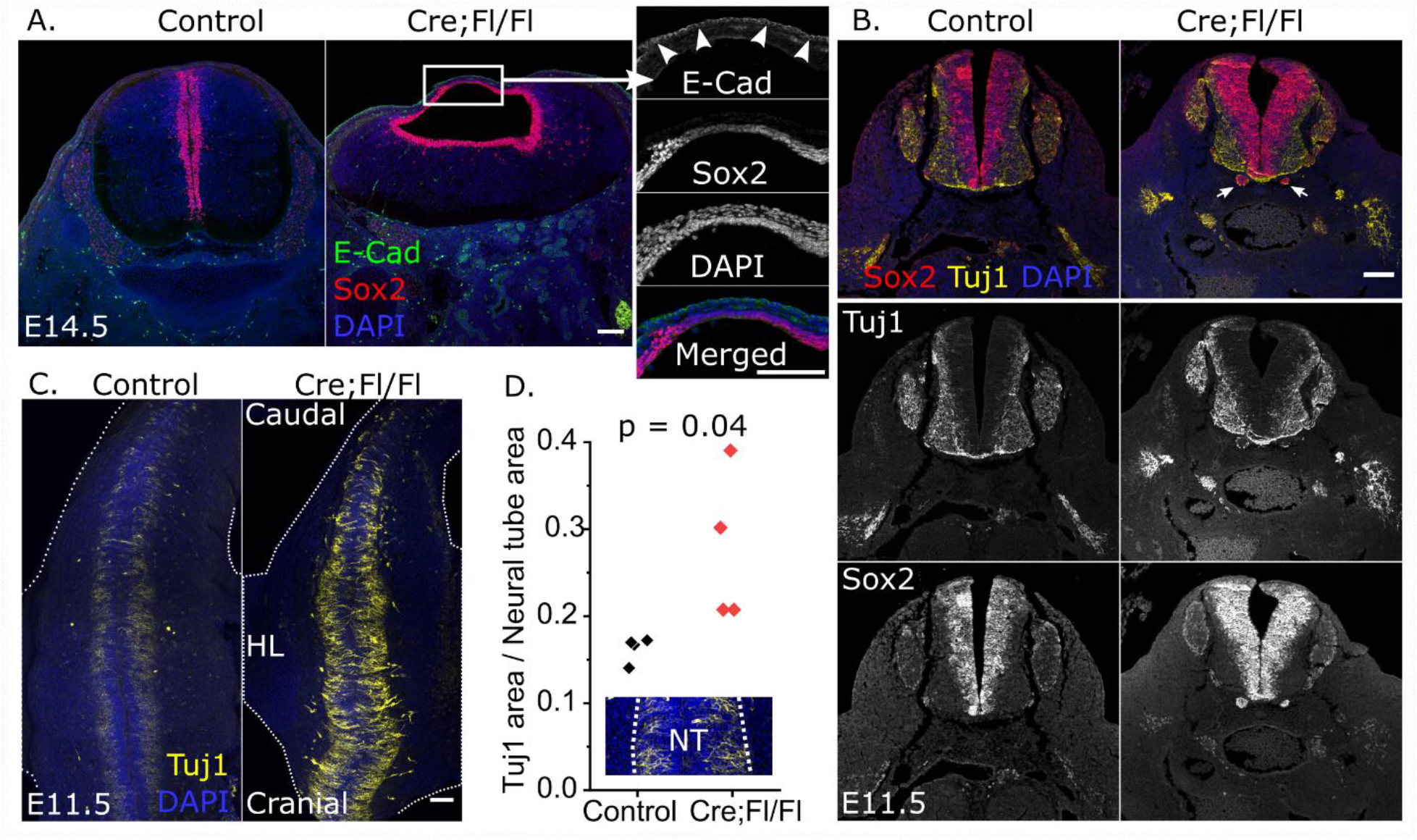
Neuroepithelial commitment to post-mitotic neurons is not diminished by loss of *Fgfr1*. A. Immunofluorescent localization of the neuroepithelial marker Sox2 and epidermal marker E-cadherin (arrowheads in insert) through the distal spinal cord of a control and Cre;Fl/Fl fetus. Scale bars = 100 μm. B. Progenitor (Sox2) and committed neuron (Tuj1) immunofluorescence of sections through the low-lumbar spinal cord of a control and Cre;Fl/Fl embryo collected at E11.5. Note ectopic clusters of Sox2-positive cells below the neural tube of the Cre;Fl/Fl embryo (white arrows, described below). Scale bar = 100 μm. C. Wholemount confocal images showing Tuj1 immunolocalization in the dorsal aspect of a control and littermate Cre;Fl/Fl embryo collected at E11.5 and imaged equivalently. Dashed white lines indicate the body outline, HL hindlimb. Scale bar = 100 μm. D. Quantification of the dorsally-visible (as in C) Tuj1-stained area as a proportion of neural tube area (NT, indicated by the dashed line in the insert).

### Regional neural tube dorsalisation precedes cystic dilation

FGF signalling has previously been found to maintain neuroepithelial cells in a progenitor state while reducing their terminal differentiation (18). Cross-sections through the lumbar neural tube of Cre;Fl/Fl embryos at E11.5 reveal distinct ventral (Shh, FoxA2, Olig2: Supplementary Figure 6A,B) and dorsal (Pax3: Supplementary Figure 6C) progenitor domains within the neural tube. Migrating neural crest cells, labelled with Sox9, can readily be identified in streams lateral to the neural tube of Cre;Fl/Fl embryos (Supplementary Figure 6C). Pax3 also immunolabels paraxial mesoderm (dermomyotome), which is markedly smaller in *Fgfr1*-disrupted embryos (Supplementary Figure 6C).

*Fgfr1* disruption causes progressive abnormalities of spinal progenitor domains. Cre;Fl/Fl embryos commonly have ectopic clusters of cells ventral to and distinct from the neural tube. These clusters express Sox2 (Figure 5B), confirming their neuroepithelial identity, as well as Shh (Figure 6A), FoxA2 (Supplementary Figure 6B), and appear to produce Tuj1-positive axonal projections (Figure 6A). They are identifiable in a short region of the neural tube between the hindlimbs and may act as an ectopic source of Shh when present alongside the floorplate and notochord (Supplementary Figure 6D). However, Shh-expressing ectopic clusters are also observable adjacent to locations devoid of detectable Shh in the ventral neural tube (Figure 6A).

**Figure 6:**
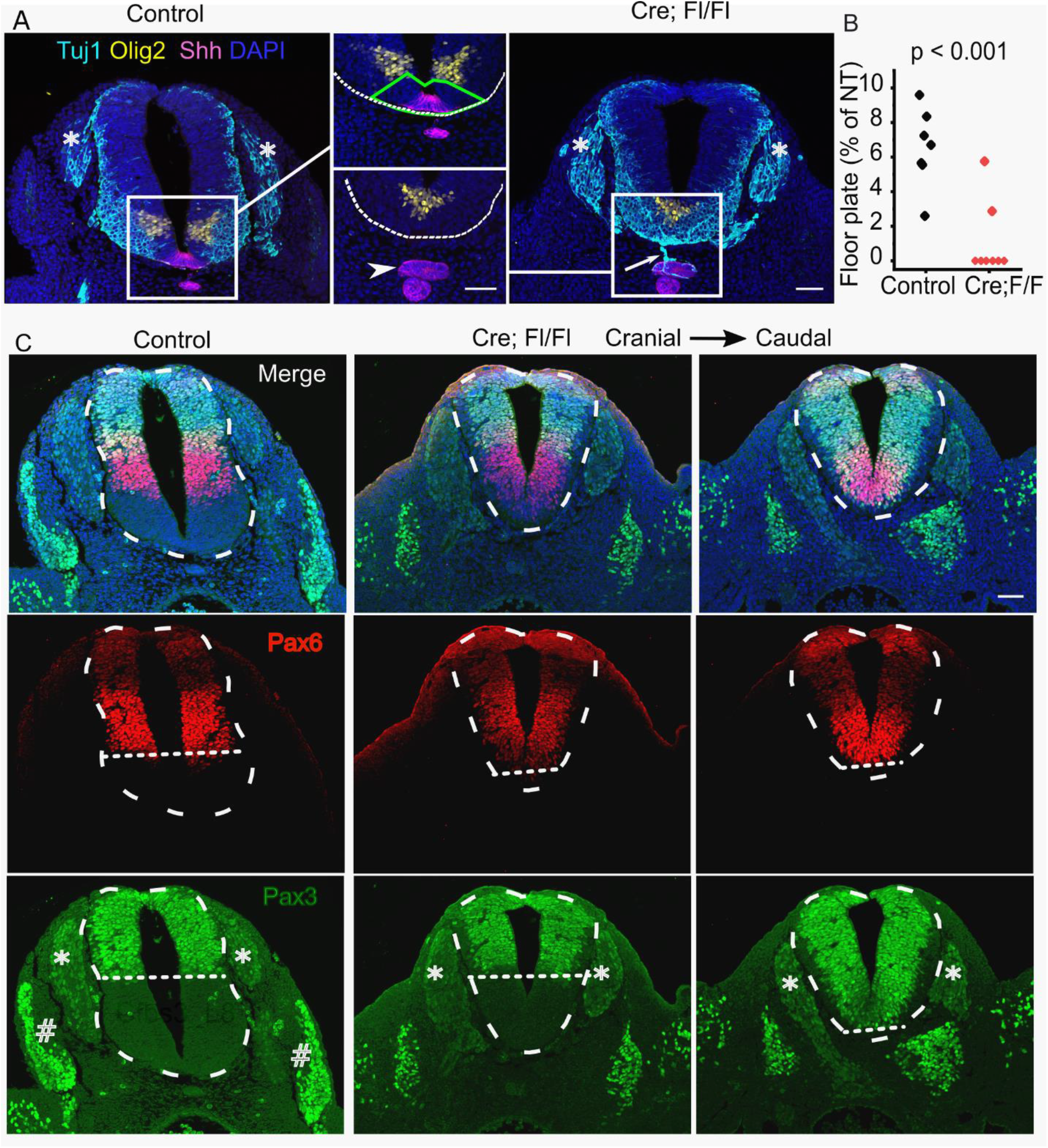
Localized, progressive loss of ventral spinal progenitor domains precedes cystic dilation of the neural tube central canal in Fgfr1-disrupted embryos. A. Immunofluorescent localization of the pMN marker Olig2, floorplate marker Shh and neuron marker Tuj1 through the lumbar spinal cord of a control and Cre;Fl/Fl littermate collected at E11. White boxes indicate the regions in the insert, dashed white lines border the ventral neural tube, green polygon indicated the region ventral to the Olig2 domain quantified in B, * indicate dorsal root ganglia, arrowhead indicates ectopic Shh-expressing tissue, arrow indicates Tuj1-positive projection from the ectopic tissue. Scale bars = 50 μm. B. Quantification of the portion of the neural tube ventral to the Olig2 domain (‘floor plate’) as a proportion of neural tube area in control and Cre;Fl/Fl embryos collected at E10.5 and sectioned between the hindlimb buds. C. Immunofluorescent localization of the intermediate neural progenitor marker Pax6 and dorsal/neural crest marker Pax3 in E11.5 embryos. Note Pax3 also labels cells in the neural crest-derived dorsal root ganglia (*) and somite-derived dermomyotome (#). Dashed white lines outline the neural tube, dotted lines indicate the ventral extent of each domain. Serial sections in the Cre;Fl/Fl embryos are ∼50 μm apart. Scale bar = 50 μm.

Regional loss of floorplate markers is observable in Fgfr1-disrupted embryos as early as E10.5 (Figure 6B), initially replaced by ventral extension of the Olig2 domain between the hindlimb buds (Supplementary Figure 6E). By E11.5, even the Olig2 domain is progressively lost as the intermediate Pax6 domain expands ventrally (Figure 6C). At more caudal levels within the same embryo, the dorsal Pax3 domain extends along the entire dorso-ventral axis of the NT (Figure 6C). Pax3 immunolabels the entire NT as it becomes abnormally circular in shape more caudally (Supplementary Figure 7A). Thus, ectopic ventral neuroepithelial clusters and progressive loss of ventral neural progenitor identities precedes NT dysmorphology which produces a terminal myelocystocele-like cystic dilation of the lumen predictably at the terminal end of the embryo (Supplementary Figure 7B).

## Discussion

The embryological origins of closed NTD sub-types have largely been inferred from their post-natal morphology and location. Animal models have been invaluable in identifying genetic/teratogenic causes of open NTDs (38), testing putative preventative agents (39), and even paving the way for spina bifida fetal surgery (40), but such models are lacking for most cases of closed NTDs. Here, we show that the human closing PNP is morphologically analogous to that of mouse embryos, and provide brightfield images suggesting it similarly adopts an elliptical structure with terminal Closure 5 at late stages of closure. The seemingly linear elevation of the caudal PNP neural folds with advancing somite stage suggests Closure 5 formation is a coordinated shape change starting at least a day earlier in mouse development.

We had previously reported impaired caudal neural fold elevation and abnormal Closure 5 associated with a caudal tissue ‘bulge’ in a model of conditional *Vangl2* deletion (6). We now observe a more exophytic outgrowth of caudal neuroepithelium where Closure 5 should have formed in embryos with pharmacologically-inhibited FGF signalling. While we do not provide a definitive explanation for this outgrowth phenotype, it may involve overactivation of retinoic acid signalling, compensatory up-regulation of pathway components as indicated by increased *Fgf8* in antagonist-treated embryos, and/or altered interactions between the neuroepithelium and paraxial mesoderm. Post gastrulation, mesoderm and neuroepithelium share a *T*-expressing neuromesodermal progenitor population. FGF signalling has been suggested to maintain and direct differentiation of this progenitor pool (41, 42). However, we do not observe gross cessation of contributions from this lineage to either neuroepithelial or mesodermal tissues within a 24-hour period of Fgfr antagonism in whole embryo culture.

Neuromesodermal progenitors contribute to axial elongation (43), for which FGF signalling is critically required (44). This is consistent with our observation of caudal truncation in embryos with caudally deleted *Fgfr1*, suggesting a non-redundant role for this receptor in axial elongation beyond the hindlimbs. Truncal elongation is unaffected despite being within the recombination domain of the Cre-driver used, suggesting receptor redundancy or activation of compensatory mechanisms which are well-described for this complex pathway (45). The concordance between failure of caudal PNP elevation and lack of Closure 5 formation seen with pharmacological antagonism of FGF receptors and selective genetic deletion of *Fgfr1* corroborates our hypothesis that completion of PNP closure selectively requires this receptor. The highly corrugated and pocketed neuroepithelium at late stages of PNP closure might produce the clusters of neuroepithelial cells observed in cross-section after the neural tube has closed in our *Fgfr1*-deletion model. Identifying the specific cell types in which *Fgfr1* signalling promotes neural fold elevation will require future tissue-specific deletion studies. Previous studies which deleted *Fgfr1* using *T*^*Cre*^ reported musculoskeletal phenotypes comparable to those observed here, but not PNP dysmorphology or later terminal myelocystocele-like phenotypes (32, 46). It is not clear whether neural tube phenotypes were carefully assessed in those studies.

The ability of embryos with caudally deleted *Fgfr1* to achieve delayed yet complete PNP closure is surprising, not only because of the level of dysmorphology their closure process overcomes, but also because alternative disruptions of the same gene cause open spina bifida in mice. Chimeric embryos containing *Fgfr1*-null cells develop spina bifida in a proportion of cases (33). Embryos in which exon 3 of *Fgfr1* is globally deleted, preventing expression of the *Fgfr1α* isoform, also develop fully penetrant spina bifida, confounded by severe dysmorphology and early lethality (47). In the current study we delete exons 8-14 (37), which include the transmembrane domain common to all isoforms (48), yet only produce closed spinal lesions at late stages of development. The open spinal neural tube we observed in a subset of embryos at E11.5 suggests predisposition to open spina bifida in our model which is not manifest in the environment and genetic background tested. Shared genetic basis between open and closed NTDs has previously been reported for exencephaly and encephalocele (49).

Closed NTD-like phenotypes have been reported in a few other models. A small, localised dilation of the closed neural tube at the level of the hindlimb buds, but not terminal myelocystocele-like phenotypes at later stages, was previously observed in *Fgf3* knockout embryos (50). At the opposite extreme, a more extensive dilation of the spinal cord’s central canal extending caudally from the thoracic spine has been described in *Noggin* knockout fetuses (51). Noggin is an antagonist of BMP signalling, which promotes dorsal spinal progenitor fates. Evidence of dorsoventral patterning disruption in *Noggin*^*-/-*^ embryos includes ventral expansion of Pax3 (52). Another striking similarity between *Noggin*^*-/-*^ and *Fgfr1*-disrupted embryos in the current study is the preferential loss of dorsoventral patterning at the level of the hindlimb buds, although *Noggin*^*-/-*^ embryos also have a markedly smaller neural tubes making direct comparisons difficult (52). One possible explanation for this regional phenotype is interactions with adjacent mesodermal structures, which are particularly abnormal caudally in our model. This is consistent with lack of sclerotome-derived vertebral bodies below the cystic lesion of *Fgfr1*-disrupted fetuses. Sclerotome has been suggested to act both as a conduit for Shh diffusion as well as being a target tissue for its action (53) and somite-derived retinoic acid is mutually antagonistic with FGF signalling in regulating caudal neurogenesis (16).

The transition from neurulation to subsequent neurogenesis continues to be explored. Premature neuronal differentiation and expansion of ventral spinal progenitor domains have been associated with open NTDs (22, 24). One model of ventral progenitor domain expansion, deletion of the mitochondrial protein *Fkbp8*, produces extensive cystic dilation of the neural tube along the entire thoracic and lumbar spine (54). Histological sections of these *Fkbp8*-deficient fetuses with ventralised neural tubes suggests thinning of the ventral spinal cord, whereas the dorsalised neural tubes of our *Fgfr1*-disrupted embryos become thinned dorsally. The mechanisms for these changes in shape are unknown, but we propose that they cause characteristic spinal dysmorphology in conditions such as terminal myelocystocele. Possible mechanisms for dorsal neural tube thinning include de-repression of differentiation causing excessive neuronal commitment and neural crest emigration, depleting dorsal progenitor populations.

The multiple interacting tissues and ubiquitous roles of FGF signalling present challenges in dissecting the molecular mechanisms by which terminal myelocystocele-like lesions emerge after closure of the neural tube. Here, we characterise a new model to begin addressing these questions. Dysmorphic PNP closure, ectopic neuroepithelial cluster formation which may act as aberrant Shh sources, enhanced neuronal production, progressive neural tube dorsalisation, and hypoplasia of surrounding mesodermal structures, all precede trumpet-like flaring of the neural tube central canal in this model. We propose that whereas failure of neurulation causes open NTDs, abnormalities in neural patterning contribute to the spectrum of clinically recognised closed NTDs.

## Materials and methods

### Animal procedures

Studies were performed under the regulation of the UK Animals (Scientific Procedures) Act 1986 and the National Centre for the 3Rs’ Responsibility in the Use of Animals for Medical Research (2019). C57BL/6 mice were bred in house and used as plug stock from 8 weeks of age. Mice were mated overnight, and the next morning a plug was found and considered E0.5. Alternatively, mice were mated for a few hours during the day, and the following midnight was considered E0.5. Pregnant females were sacrificed between E9.5 and E16.5.

*Fgfr1*^*Fl/+*^ mice were as previously described (37). *Cdx2*^*Cre/+*^ mice (36) were used to breed *Cdx2*^*Cre*^*Fgfr1*^*Fl/+*^ stud males. The stud males were then crossed with *Fgfr1*^*Fl/Fl*^ females to obtain *Cdx2*^*Cre*^*Fgfr1*^*Fl/Fl*^ embryos. *Cdx2*^*Cre*^*Fgfr1*^Fl/+^ and Cre-negative embryos were phenotypically normal and considered littermate controls. *Rosa26-mTmG* reporter mice were as previously described (55). For lineage tracing, *Cdx2*^*Cre*^*Fgfr1*^*Fl/+*^ stud males were crossed with *Fgfr1*^*Fl/Fl*^*mTmG* females.

To lineage-trace neuromesodermal progenitors during Fgfr pharmacological inhibition, *T*^*CreERT2/+*^ stud males (56) were crossed with *mTmG* females and tamoxifen (10 mg/mouse) was administered orally at 6 pm on E8.5 as previously validated (57). Embryos were then explanted into whole embryo culture starting at E9.5 and fixed 24 hours after the start of culture.

The RARE-hsp68LacZ reporter mice (JAX stock #008477) (58) were on a CD1 background as previously described (58).

### Human embryos

One CS11 and one CS13 human embryo collected contemporaneously with this project were provided by the Human Developmental Biology Resource tissue bank. These embryos were non-destructively imaged using reflection confocal microscopy and returned to the tissue bank. Brightfield images of CD9-CS12 embryos collected by the tissue bank were also provided.

### Embryo culture

Whole mouse embryo culture was performed in neat rat serum following published protocols (59). Pharmacological inhibitors pan-FGFRi (BGJ 398, Generon, used at 1 μM) or FGFR1-targeting inhibitor (PD 173074, Cambridge Bioscience, used at 1 μM) were dissolved in DMSO and added to the rat serum at the start of culture. For control conditions 0.1% DMSO was added to the rat serum. Embryos were size matched and randomized to inhibitor or vehicle groups using a coin toss.

### Immunofluorescence

Embryos were dissected from their extraembryonic membranes, rinsed in ice cold PBS and fixed in 4% PFA overnight. Whole-embryo immunostaining was as previously described (5). Primary antibodies were used in 1:100 dilution and were as follows: rabbit E-cadherin (3195, Cell Signaling Technology), mouse N-cadherin (14215S, Cell Signaling Technology), rabbit pERK (#9101, Cell Signalling Technology), mouse Fgfr1 (ab824, Abcam).

For N-cadherin staining, antigen retrieval was first performed for 25 mins at 100°C using 10 mM sodium citrate with 0.05% Tween 20, pH 6.0. For pERK staining, a methanol/acetone post-fix step was added after PFA fixation. Embryos were washed with ice cold 50:50 methanol/acetone for 30 min, followed by decreasing concentrations of methanol (10 min each) prior to blocking in 5% BSA solution in PBS with 0.1% Triton X-100. Secondary antibodies were used in 1:200 dilution and were Alexa Fluor conjugated (Thermo Fisher Scientific). Alexa Fluor 568 or 647– conjugated Phalloidin was from Thermo Fisher Scientific.

For immunostaining of paraffin sections, antigen retrieval was always performed prior to blocking in 5% BSA solution. Primary antibodies were applied overnight at 4°C (1:100-1:200 dilution). Primary antibodies were as follows: rabbit anti-Sox2 (ab97959, Abcam), rabbit anti-Sox9 (7H13L8, Thermo Fisher Scientific), mouse anti-Tuj1 (MMS-435P, BioLegend), goat anti-Olig2 (AF2418, Novus Biologicals), rabbit anti-Shh (C9C5, Cell Signalling Technology), mouse Pax3 (Pax3-c, DSHB), rabbit anti-Pax6 (901301, BioLegend), goat anti-FoxA2 (AF2400, R&D Systems). Sections were washed, incubated with the secondary antibodies above (1:500 dilution) and counterstained with DAPI for 2 h at room temperature.

### Confocal microscopy and image analysis

Images were captured on a Zeiss Examiner LSM 880 confocal using 10 ×/NA 0.5 or 20 ×/NA 1.0 Plan Apochromat water immersion objectives. Images were processed with Zen 2.3 software and visualized as maximum projections in Fiji (60). Inset in Figure 1F was obtained with AiryFast scanning mode using the 20×/NA 1.0 water immersion objective. Reflection images of CS11 and CS13 human embryos were captured at 633 nm wavelength using a 10×/NA 0.5 water immersion objective. The z stacks were 3D rotated and visualised as maximum projections. Sections were captured with AiryFast scanning mode using the 10×/NA 0.5 dipping objective. Salt and pepper noise was removed when appropriate using the de-speckle or ‘remove outliers’ function in Fiji.

For morphometric analysis, PNP length and width were calculated by annotating the PNP rim and then measuring the major and minor axis using the fit ellipse function in Fiji. Neural fold elevation was measured in optical reslices of confocal Z-stacks. Distances between the limb buds and the caudal end of the embryos were measured in stereoscope images using the segmented line tool in Fiji. Floor plate area was measured using the polygon tool and is expressed as percentage of total neural tube area.

### Histology and X-gal staining

Following overnight fixation in 4% PFA, E11.5 embryos were dehydrated with increasing concentrations of ethanol (20 min each). They were then moved to 100% histoclear for 20 min at room temperature, washed twice with 50:50 histoclear/paraffin mix for 10 min at 65°C and then embedded in 100% paraffin. Paraffin was replaced three times to ensure removal of histoclear. Sections were made at 10 μm thickness. The protocol was similar for E14.5 fetuses except for longer dehydration steps (40 min each), followed by overnight storage in 100% ethanol. In this case, histoclear was added for 40 min at RT. Sections were either antibody stained (see immunofluorescence) or stained with hematoxylin and eosin. For bone staining, P1 pups were fixed overnight in 100% ethanol followed by three days in 100% acetone. They were then stained with Dawson staining solution overnight at 37°C, washed and cleared with 1% KOH at 37^°^C. Slower clearing was achieved with 10% glycerol in 1% KOH at RT. For Xgal staining, embryos were removed from culture and fixed in 2% PFA for 10 min, rinsed in 2 mM MgCl_2_in PBS and incubated in staining solution: 1× PBS, 20mM K3Fe(CN)6, 20mM K4Fe(CN)6-3H2O, 2mM MgCl2, 0.01% DOC (=sodium deoxycholate), 0.02% NP-40 with X-gal (1 mg/ml final concentration) as a 1:50 addition at 37°C for 3 hours.

### In situ hybridization

In situ hybridisation of whole E10.5 embryos was performed essentially as previously described (61) using a cDNA probe for *Fgf8* ((62) *variant 4*, insert 800 bp) cloned into a pBluescript SK(+) vector. Sense and anti-sense riboprobes were generated using a digoxygenin RNA labelling kit and T3/T7 RNA polymerases (Roche). Images are representative of at least 3 embryos per condition.

### Statistical analysis

Statistical analysis was performed in OriginPro 2017 (Origin Labs). Individual embryos were the unit of measure. Comparison of two groups was by Student’s *t* test. Proportions were compared using Fisher’s exact tests. Comparisons of slopes were based on Pearson’s correlation. Graphs were made in OriginPro 2017 and are shown as scatter plots with individual embryos as the unit of measure. Analysis of Closure 5 presence was carried out blinded to genotype/treatment group, although marked PNP dysmorphology could not be masked in these analyses.

## Acknowledgements

This study was supported by the Wellcome Trust (211112/Z/18/Z and 211112/Z/18/A, both to GLG). The NIHR Great Ormond Street Hospital Biomedical Research Centre (NIHR GOSH BRC) supports research infrastructure at the UCL GOS Institute of Child Health. The views expressed are those of the authors and not necessarily those of the NHS, the NIHR or the Department of Health. Use of *Fgfr1*-floxed alleles was with permission of their creator, Prof Chu-Xia Deng, and we are grateful to Dr Emma Rawlins for providing these mice. We thank Mr Dominic Thompson for discussions of terminal myelocystocele phenotypes.

