## Supplementary for "Caudal *Fgfr1* disruption produces localised spinal mis-patterning and a terminal myelocystocele-like phenotype in mice"

### Supplementary Figures and Legends

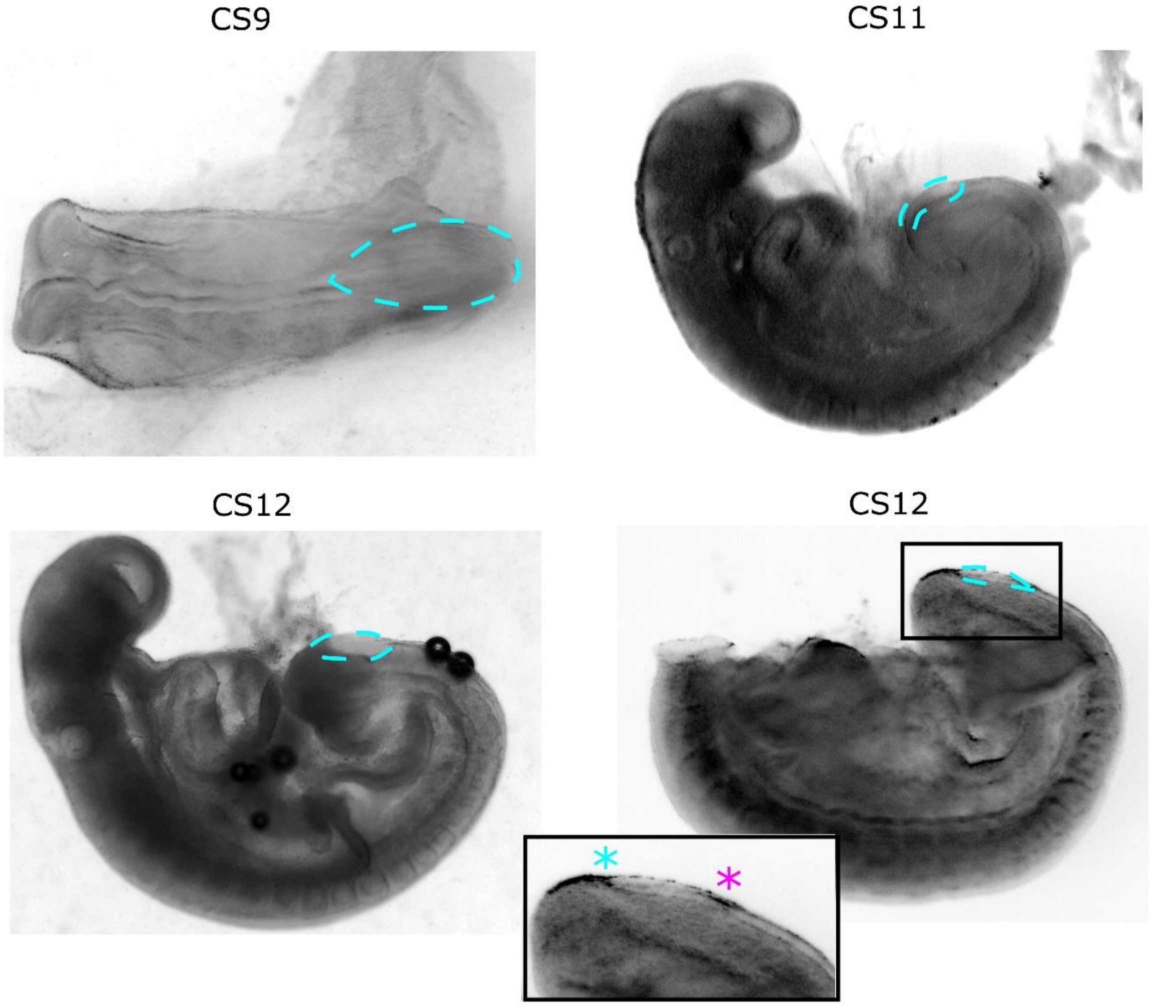

**Supplementary Figure 1: Human PNPs have an elliptical morphology suggestive of Closure 5 formation at late stages of closure.**

Brightfield images of human embryos at the Carnegie Stages (CS) indicated. The dashed cyan line annotates the posterior neuropore, which at CS9-11 has a spade-like morphology. By CS12, the human PNP has acquired an elliptical shape and the anatomy of the caudal extremity suggests formation of Closure 5 (cyan asterisk). The magenta asterisk indicates the rostral-to-caudal zippering point.

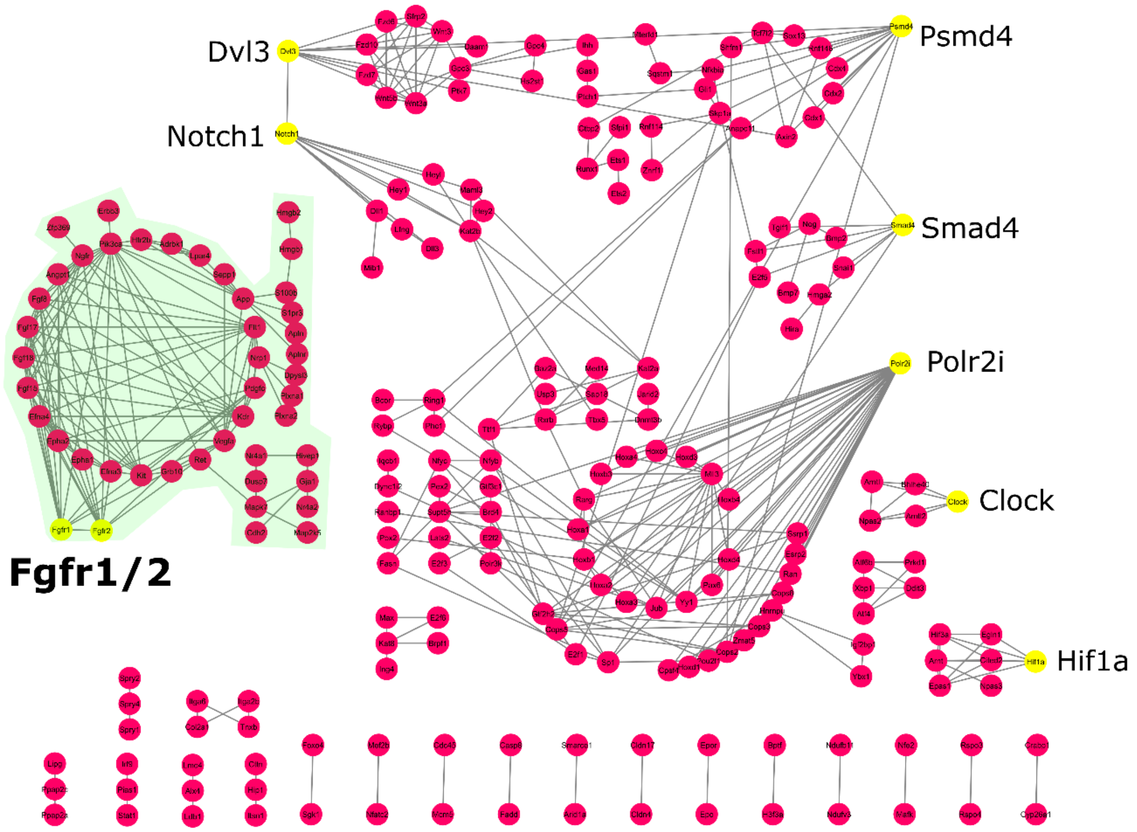

**Supplementary Figure 2: Fgf pathway components are known to be robustly expressed in the tissue region where Closure 5 forms.**

523 genes known to be expressed in the region of Closure 5 formation (tail bud) between E9.5-E10.5 in mouse embryos were identified using the EMAGE gene expression database (http://www.emouseatlas.org/emage/)(1). Genes whose protein products are known to interact with high confidence in reported experiments and curated databases were identified in StringDB(2); non-interacting members were suppressed. The resulting network of interacting Closure 5-region genes was analyzed in Cytoscape(3) to identify sub-networks linked to nodes of interest (yellow, gene names annotated). Genes interacting with FGFR1 and 2 are indicated with green shading. Yellow nodes indicate genes related to specific pathway of interest. Results were obtained from bioinformatic analysis.

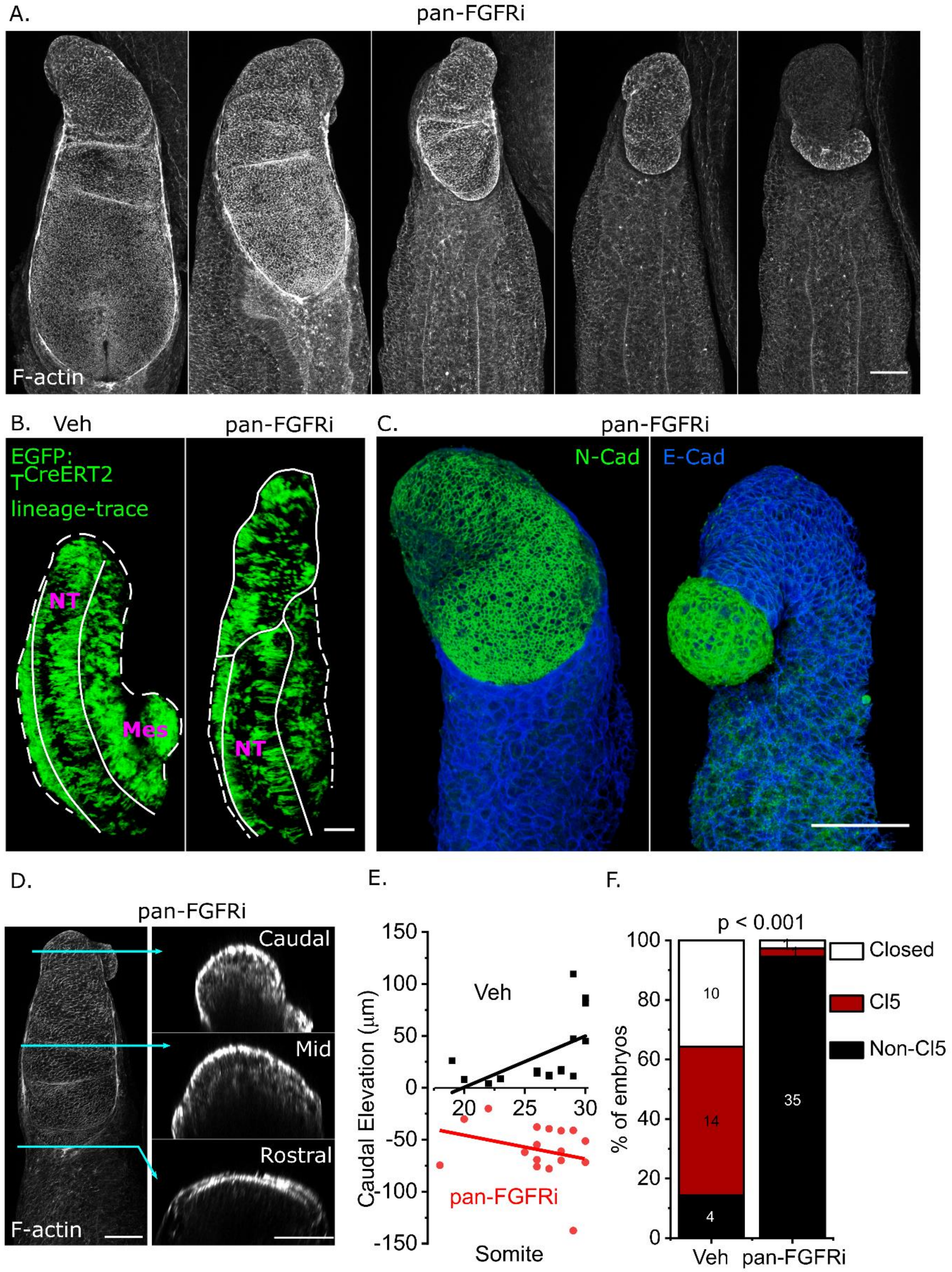

**Supplementary Figure 3: Pharmacological inhibition of Fgf signaling in cultured mouse embryos prevents neural fold elevation required for Closure 5 formation.**

A. Confocal images of phalloidin stained embryos cultured with pan-FGFR inhibitor for 24 hours.
B. 3D reconstructions of confocal-imaged vehicle and a pan-FGFR inhibitor treated embryo after 24 hours whole embryo culture. Neuromesodermal progenitors are lineage traced with $T^{CreERT2}$ and continue to give rise to both neural tube (NT) and mesoderm (Mes) in both conditions.
C. 3D reconstructions of confocal-imaged embryos after 24 hour culture with pan-FGFR inhibitor. Immunolocalisation of E- and N-cadherin shows that the PNP overgrowth is neuroepithelial.
D. Confocal image of a phalloidin stained embryo after 24 hours culture with pan-FGFR inhibitor. The cyan arrows indicate the level of the optical cross-sections at the rostral, mid and caudal level.
E. Quantification of elevation at the caudal neural folds (90% of the PNP's length) in embryos at the indicated somite stages.
F. Quantification of the proportion of embryos collected at E10.5 with closed PNPs, open PNPs with Closure 5 morphology or open PNPs without Closure 5. Numbers indicate the number of embryos observed in each category.

Scale bars = 100 µm.

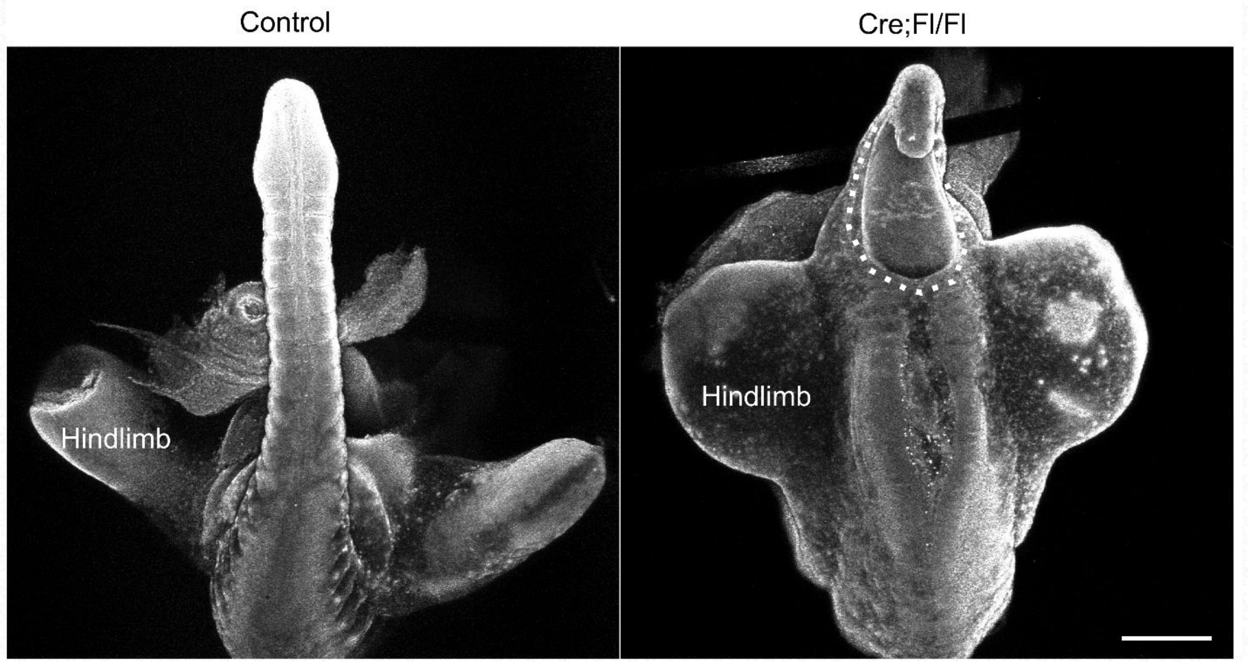

**Supplementary Figure 4: A subset of Fgfr1-disrupted embryos have an open distal spinal lesion as late as E11.5.**

Reflection images of control and Cre;Fl/Fl littermates collected at E11.5. The dotted white line encircles the open lesion at the caudal end of the body axis. Scale bar = 500 µm.

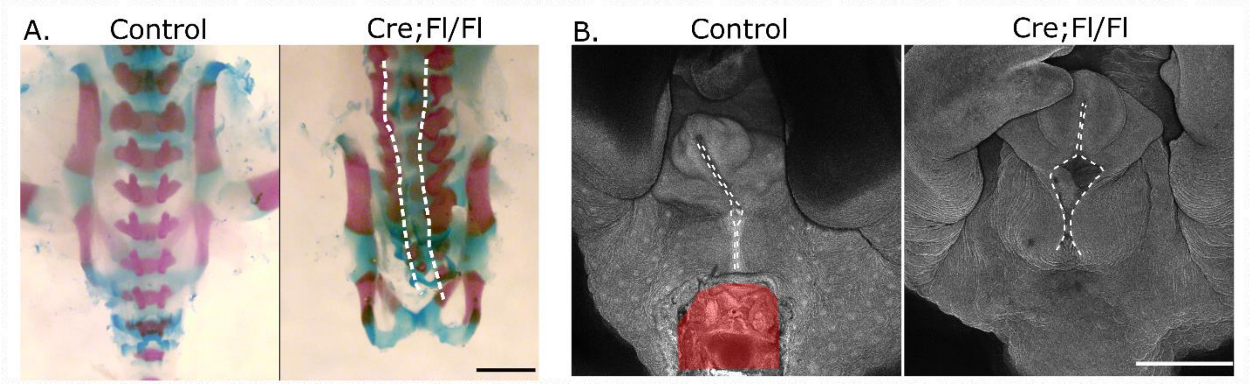

**Supplementary Figure 5: Skeletal and perineal abnormalities in embryos with caudally-deleted *Fgfr1*.**

A. Whole-mount skeletal staining with Alcian blue and Alizarin red of control and Cre;Fl/Fl P1 pups from the same litter. Dashed white lines indicate bifid neural arches. Scale bar = 850 µm.
B. Reflection images of control and Cre;Fl/Fl littermates collected at E16.5. The dotted line annotates the perineum, which is fused in control and partially open in the mutant. Red area shows the tail base, which was cut to enable imaging of underlying structures in the control fetus. Scale bar = 1 mm.

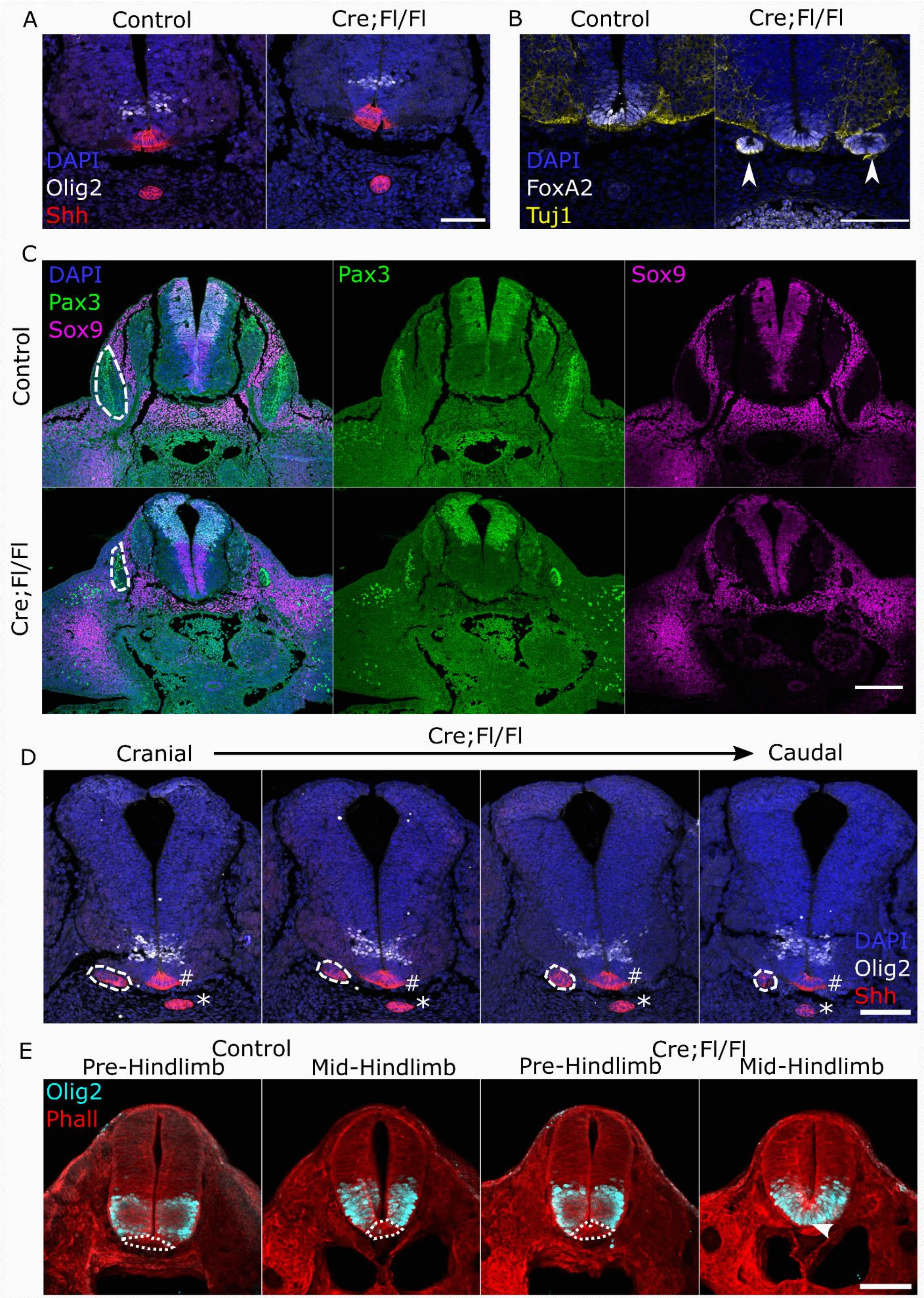

A
Control
Cre;Fl/Fl
DAPI
Olig2
Shh
B
Control
Cre;Fl/Fl
DAPI
FoxA2
Tuj1
C
DAPI
Pax3
Sox9
Pax3
Sox9
Control
Cre;Fl/Fl
D
Cranial
Cre;Fl/Fl
Caudal
#
*
DAPI
Olig2
Shh
E
Control
Cre;Fl/Fl
Pre-Hindlimb
Mid-Hindlimb
Pre-Hindlimb
Mid-Hindlimb
Olig2
Phall

**Supplementary Figure 6: Caudal *Fgfr1* deletion causes localized progenitor domain abnormalities.**

A-C Immunofluorescent localization of neuronal progenitor markers through the lumbar spinal cord of control and Cre;Fl/Fl littermates collected at E11 showing: A) pMN marker Olig2 and floor plate and notochord marker Shh. B) Post-mitotic neuronal marker Tuj1 and floor plate marker Foxa2. The arrowheads indicate ectopic Foxa2 foci in the Cre;Fl/Fl embryo. C) Dorsal/neural crest marker Pax3 and neuroepithelial/neural crest marker Sox9. The dashed line encircles the dermomyotome which is markedly reduced in the Cre;Fl/Fl embryo. Scale bars A,B = 100 µm, C = 200 µm.

D. Serial sections through the lumbar spinal cord of a Cre;Fl/Fl embryo collected at E11 and stained for Olig2 and Shh. Shh stains the floor plate (#), notochord (*) and ventral ectopic clusters encircled by a dotted line. Scale bar = 100 µm.

E. Sections at pre- and mid-hindlimb level of control and Cre;Fl/Fl littermates collected at E10.5. The Olig2 negative floor plate (dotted line), disappears at the mid-hindlimb level of the mutant (white arrowhead). Scale bar: Scale bar = 100 µm.

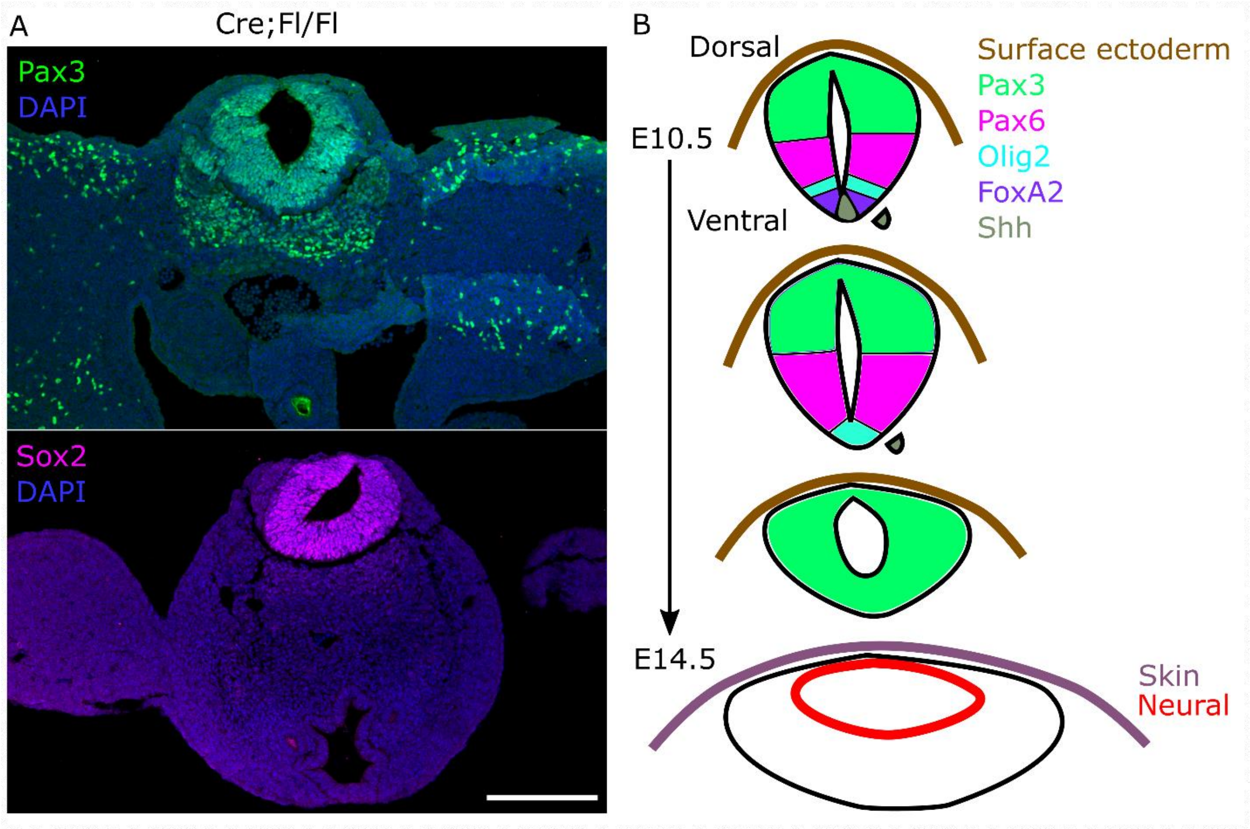

**Supplementary Figure 7: Neural tube dorsalization precedes its dysmorphology and central canal dilation in Fgfr1-disrupted embryos.**

A. Hindlimb level section through a Cre;Fl/Fl embryo collected at E11.5. The neural tube lumen is circular and stains positive for box Sox2 and Pax3 throughout. Scale bar = 200 µm.

B. Schematic representation showing the progressive neural tube dorsalisation after caudal deletion of *Fgfr1* in Cre;Fl/Fl embryos. Ectopic clusters of Shh and Foxa2 at E10.5 precede loss of ventral progenitor domains. This leads to ventral expansion of Pax3 and Pax6 from E11.5, which finally encircle the neural tube lumen. Central canal dilation produces a terminal myelocystocele-like phenotype at E14.5.
